# Universe of Lasso Proteins: Exploring the limit of entanglement and folding landscape of proteins predicted by AlphaFold

**DOI:** 10.1101/2025.03.21.644650

**Authors:** Fernando Bruno da Silva, Agata P. Perlinska, Jacek Płonka, Erica Flapan, Joanna I. Sulkowska

## Abstract

Knots and lasso topology represent a class of natural motifs found in proteins which are characterized by a threaded structure. Proteins with a lasso motif represent a macroscopic version of the peptide lasso, which are known for their high stability and offer tremendous potential for the development of novel therapeutics. Here, based on AlphaFold, we have shown the limit of topological complexity of naturally occurring protein structures with cysteine bridges. Based on 176 million high confidence (pLDDT > 70) AlphaFold-predicted protein models and a detailed analysis of the conservation of the motif in a family, we found four new lasso motifs, including L_4_ and LS_4_LS_3_ topologies, and the first examples of knotted lasso proteins: L_1_K3_1_ and L_3_#K3_1_. We show that in the case of natural proteins, there are no lassos with 5 threadings but there exist some with 6. Families possessing proteins with more than 6 threadings did not exceed the conservation threshold of 10%. Moreover, we propose a probable folding mechanism for the LS_4_LS_3_ lasso motif, enhancing our view on protein folding and stability. This work expands the topological space of lasso type motifs in proteins but also suggests that more complex structures could be unfavorable for proteins.

**Highlights:**

- Discovery of novel non-trivial lasso motifs: the L_4_, supercoiling of both tails LS_4_LS_3_, and the first knotted lasso proteins: L_1_K3_1_ and L_3_#K3_1_.
- The knotted lasso motifs are in membrane proteins.
- Lassos topologies with 5 or more crossings are not conserved in protein families, and more complex motifs do not exist
- 472 new InterPro entries with a high probability of non-trivial lasso motif
- Potential folding pathway for proteins with complex supercoiled lasso motif LS_4_LS_3_

**Graphical Abstract:** 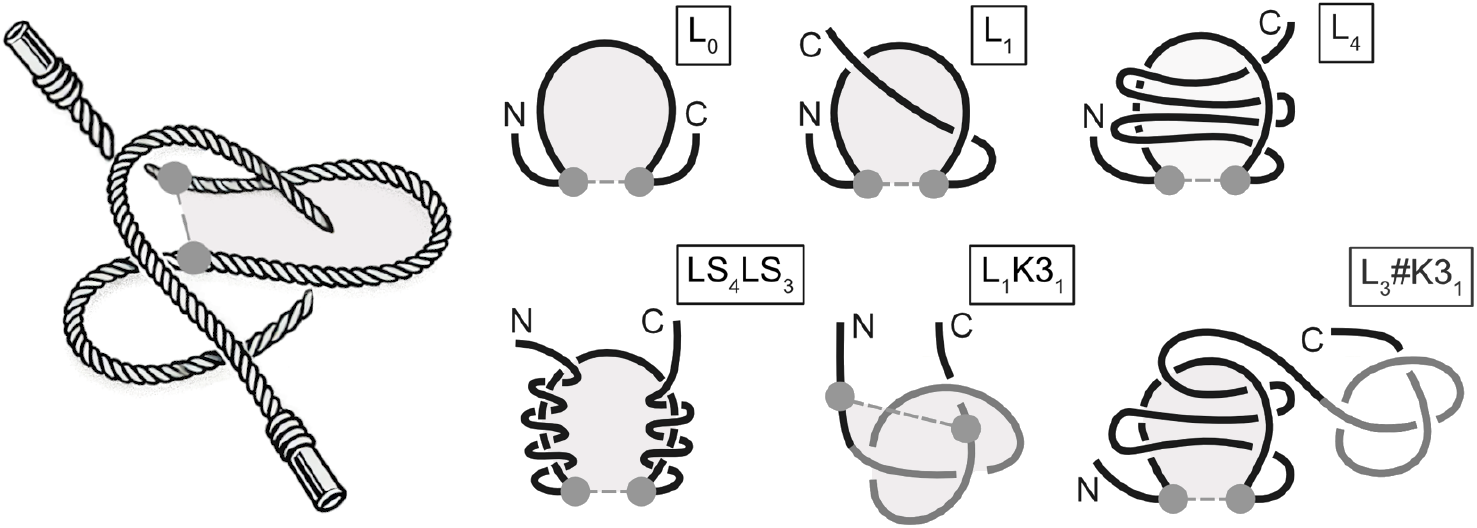

## Introduction

A lasso in the microscopic world is a concept similar to its equivalent in the macro world – a loop made on a rope. In proteins, the rope is a polypeptide chain, and the loop is made by a covalent link between two cysteine residues, the disulfide bridge. However, lassos are found not only in proteins but also in peptides [1] which form very stable structures. The peptide lassos, due to their small size, are a well-studied group of molecules with a known function and a wide range of possible applications [2, 3]. On the other hand, lassos in proteins are less known, even though they are shown to be present in several different protein families. Their size, complexity, and difficulty in assessing their function hinder experimental studies [4, 5, 6]. Here, using a theoretical approach and structural data obtained from machine learning algorithms, we characterize protein lasso motifs and show novel entangled topologies that provide insight into the complexity of protein structures towards practical applications [7].

The topic of protein entanglement is important and has started to have real-world applications. It was shown that topologi engineering in the form of the introduction of catenation holds great promise in the development of therapeutic proteins (artificial antibody with enhanced affinity and *in vivo* stability [8]). With progress in topological engineering for proteins, the lasso motif gives a new pathway in the development of therapeutic proteins [7].

Lasso topology in proteins is divided into types depending on the complexity of the motif. It is measured by the number of times the polypeptide chain pierces the lasso loop. Note that in proteins, after the lasso loop is formed, there are two parts of the remaining chain, one closer to the N- and the other to the C-terminus (N- and C-tail). Therefore, when determining the exact topology, we take into account which tail threads the loop.

The first large-scale classification of protein lasso motifs was done in 2016 in the LassoProt database [9]. In the case of one tail threading, four main motifs were found in proteins: L_1_, L_2_, L_3_, and L_6_ (note that there are lasso proteins with six piercings, but none with four or five) [10, 9]. There are also instances when both tails pierce the loop (a two-sided lasso): L_1,1_, L_1,2_, L_2,1_, and L_4_,_3_ [9, 11]. The most complex type of lasso known to date, based on proteins deposited in the PDB and LassoProt, is supercoiling when the tail is piercing and wrapping around the loop. There are three motifs with supercoiling known: LS_2_, LS_3_, and LS_4_ [11]. When none of the tails pierce the loop, the lasso is trivial and is denoted L_0_. Lasso proteins of this type are the majority; however, they are not classified as having a complex and entangled topology – they just possess a disulfide bridge.

Besides lasso, there are other types of entanglement present in protein structures [12]. Several loops closed (but not pierced) by cysteine bridges form a topologically trivial structure called a cysteine knot [13, 14]. When the loops are pierced by each other, we speak of links [15]. Structures with multiple bridges with closed loops (intersecting or not), forming a *θ* shape, are called theta curves [16]. Entanglement made solely on the polypeptide chain and not locked by cysteine bonds is called a knot. Interestingly, among all these topologies, the largest group is formed by proteins with a single-pierced lasso motif (L_1_) [6], even though they are not a well-researched group.

Based on experimentally obtained protein structures, it was shown that 39% of the lasso proteins are classified as enzymes [11]. Specifically, their function seems to depend on the type of the lasso motif. In individual cases, such as leptin, its L_1_ lasso is known to influence the activity, dynamics of the native state, and stability of the protein [17]. Recently, computational studies have explored and identified favorable folding pathways for leptin [18, 19]. However, the folding mechanisms of lasso topology proteins are still poorly understood, requiring further experimental and theoretical investigations.

As the machine learning revolution also affected structural biology, we now have several orders of magnitude more 3D structures available to explore, including structures we had not previously expected to exist. For example, AlphaFold predicted new types of knotted [20, 21, 22] and linked proteins [23], coming from either known knotted families or not. Importantly, experimental studies have already confirmed the existence of several new knotted types: double knotted 3_1_#3_1_ [24] and 7_1_ [20] – the first type of knot which is neither a twist type nor has unknotting number one. Moreover, this amazing achievement was possible without any homologous structures (the 7_1_ knot). Given AlphaFold’s capability to predict new entangled folds, we explore whether new lasso motifs are also feasible.

The topology should be conserved in the majority of the proteins in a given family, if it is important for the biological function or the environmental conditions [25, 26, 27, 28]. Along with the amount of available structural data, this provides an opportunity for both quantitative and qualitative analysis of predicted protein structures. It is particularly important for the lasso motifs, since they are less understood than knots. Specifically, it is known that a knot is perfectly conserved in a protein family [29, 22], which has not yet been studied for protein families containing lasso motifs.

Herein, we have conducted a comprehensive review of 176 million high-quality structures predicted by AlphaFold 2. For each protein structure, we determined the dominant lasso (in the AlphaLasso database [30]) and knot type (in the AlphaKnot database [31]). As a result, we found 2.5 million proteins with non-trivial lasso topology that we further analyzed. Based on this data and rigorous analysis of all the proteins in the given family, we show new types of lassos and the first knotted lasso proteins.

## Results

We based our study on all protein structures predicted by AlphaFold that possess a non-trivial topology such as a lasso or knot. We used two databases: AlphaLasso [30] and AlphaKnot [31], where all such structures are deposited. In total, we evaluated over 2.5 million high-quality (pLDDT over 70) protein models with a lasso topology. Since one protein structure can have more than one lasso motif, we identified 3.1 million lasso motifs. In the case of proteins with knots, we collected information about 700 thousand structures. We classified each protein by a family (using the InterPro database, see Methods section for details) to analyze the level of conservation of the topology within groups of similar proteins. Moreover, we used it to find new and robust lasso motifs that are preserved in a given family (Figure 1).

**Figure 1:**
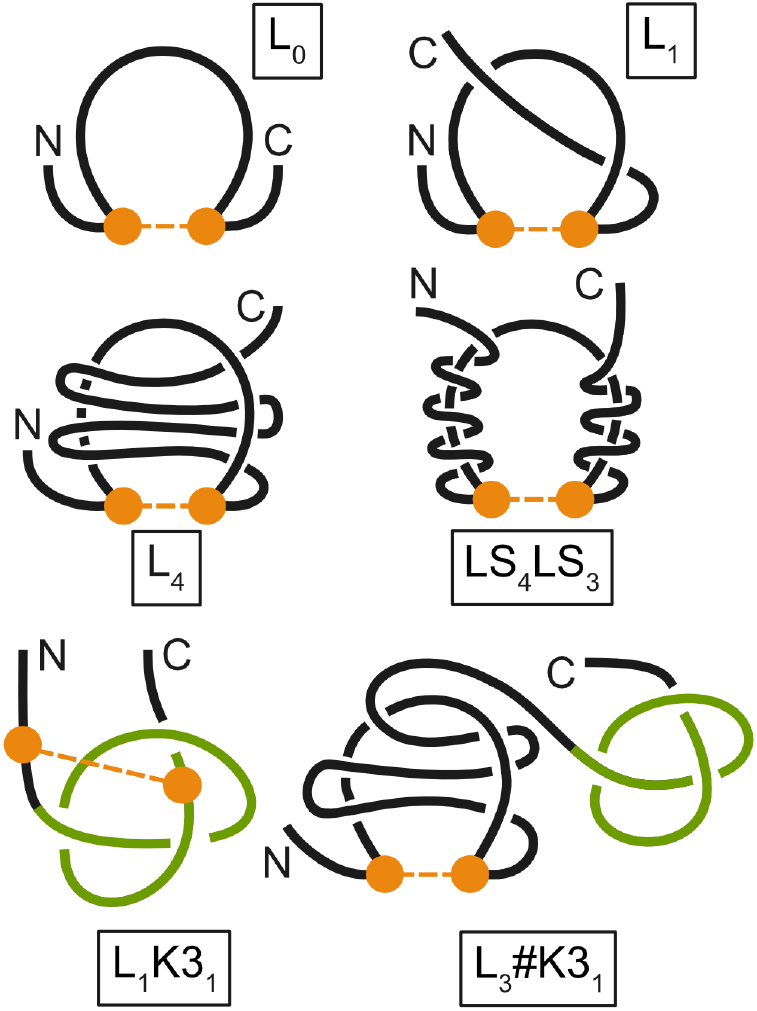
Known and new lasso types. The first row shows the simplest lasso types (L_0_ with no piercings, and L_1_ with a single piercing through the loop) that are known to be present in protein structures. The remaining are new lasso types presented in this study and found in modeled protein structures. L_4_ with a single tail piercing the loop four times, LS_4_LS_3_ with both tails piercing and wrapping around the loop (supercoil) four and three times, L_1_K3_1_ with a lasso and knot intertwined, and L_3_#K3_1_ with first a lasso loop pierced three times and next the 3_1_ knot. Orange beads and lines connecting them depict a cysteine bridge that forms the lasso loop. The knot is green-colored.

### Statistical analysis of lasso motif conservation in the family

Since it was shown that AlphaFold is capable of predicting new knot topologies in proteins [32, 20], we now ask whether the same can be done for new types of lassos and, in general, how well the lasso motif is conserved in a given family [33]. However, unlike the knotted topology, which is strictly conserved within a family [33, 29], the existence of lassos can depend on a single cysteine mutation. The second difficulty is the identification of the cysteine bridge, which is not annotated in the 3D structures provided by AlphaFold. Therefore, we use a distance cutoff for the bridge – the thiol groups have to be within 3 Å of each other.

To carry out a qualitative analysis of lasso motives provided by AlphaFold, we focused on the most prevalent protein families. Since not every protein has an AlphaFold-generated model, some of the families have less structural information than others. Given that the probability of correctness of the new topology is higher, the better its conservation in the family, we opted for the families for which we had sufficient topological information – meaning at least 30% of the family members were analyzed by the AlphaLasso web server. 1041 families fulfilled these criteria. We then categorized the families by lasso topology, excluding L_0_. On this basis, we selected only 6 motives, *L*_1_, *L*_2_, *L*_3_, *L*_4_, *L*_6_ (no *L*_5_), out of the 24 possible ones that the AF predicted.

First, we analyzed the level of conservation of the lasso motif in protein groups (based on InterPro) that have at least a single experimental confirmation of the motif (Figure 2).

**Figure 2:**
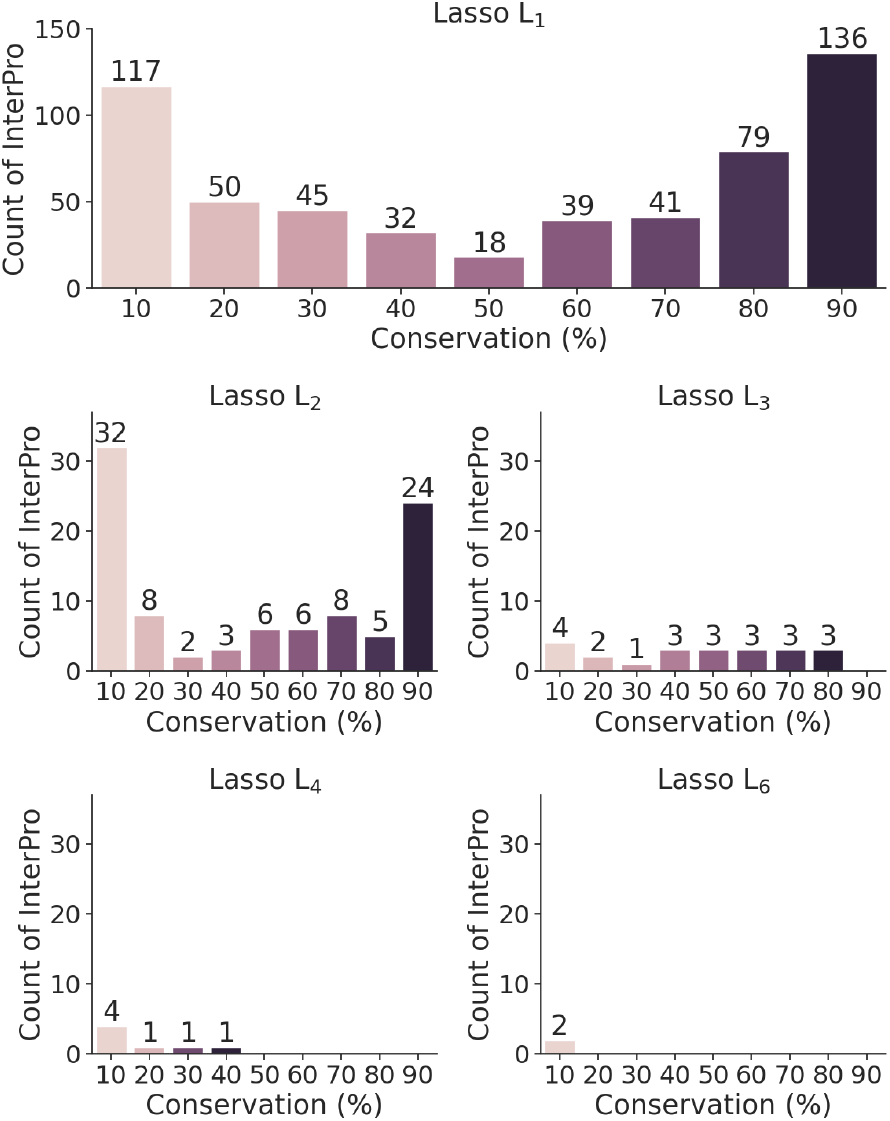
Conservation level of the main lasso types in known entangled protein groups. Shown is the number of InterPro entries possessing at least one experimentally confirmed lasso structure. The conservation level is calculated as the number of proteins exhibiting a given topology to the number of analyzed proteins in this InterPro entry and presented as bins (e.g. conservation of 10% covers values in range 10%-19%). Only entries where at least 50% (minimum 100) of all proteins present in the InterPro entry were analyzed and with conservation at least 10% are considered.

The results show a great variety – we encountered virtually every conservation level. However, for the L_1_ and L_2_ lasso types, there are two leading values: 10-20% that show no conservation, and more than 90% that show strict conservation of the lasso. Therefore, the level of conservation of the lasso depends on the given protein group, contrary to the protein knots whose presence is strictly conserved in a family.

Comparing data from AlphaLasso to data deposited in the PDB, we found that in 569 families (InterPro entries) indeed, the topology predicted by AlphaLasso agrees with the topology of the structure deposited in the PDB. On the other hand, this means that we have found 472 new entries where the topology may be correct, and its correctness may be suggested by the motif behavior in the family.

Figure 3 also shows a clear pattern: the more complex a lasso topology, the less it is conserved. In fact, when lassos have more than six piercings, they are not conserved (present in less than 10% of proteins from a given family). During the analysis, we found proteins with very complex lasso motifs with as many as 18 intersections. This suggests that very complex lasso shapes might be harder to maintain during evolution. However, it should be noted that there may be some isolated exceptions where only one or two sequences from the whole family have such a complex topology. These data are available for further research.

**Figure 3:**
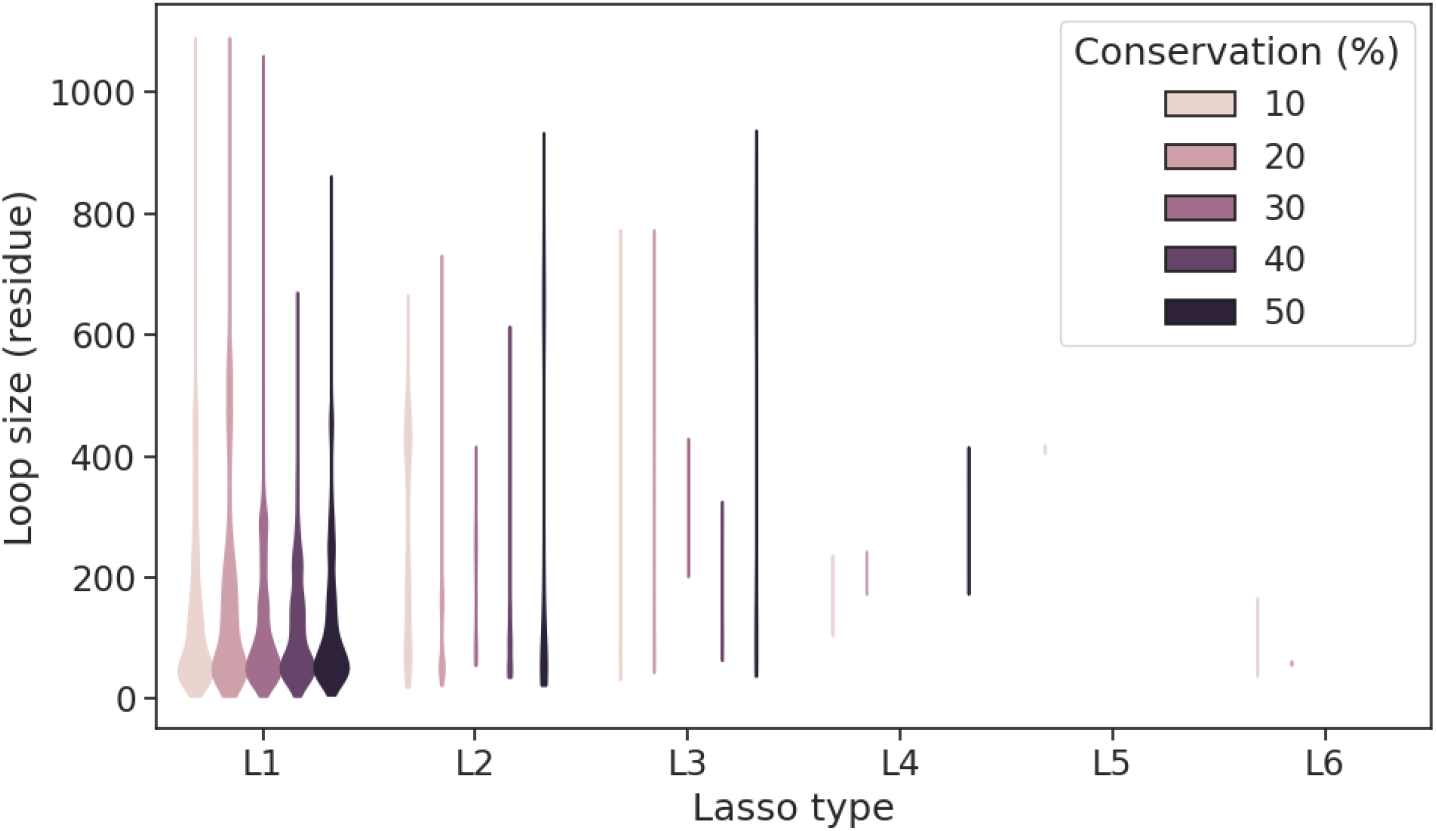
Distribution of loop size by lasso types. A distribution of loop size is shown, divided by the lasso type and its conservation across the InterPro entry. Only the entries where AlphaLasso analyzed at least 50% (minimum 100) of all proteins annotated in the InterPro database were considered. Lassos of the same type present in the same protein were grouped based on their position and an average of their loop size was taken.

Having such a verified dataset (Figure 3), we found that the most common lasso loops are small, between 5 and 100 amino acids long. These smaller loops may give proteins an advantage by making them more stable. To the best of our knowledge, due to a lack of data, this has never been shown before.

### New lasso motifs in proteins

Among 2.5 million protein structures with lassos, we identified four new motifs (Figure 1). To identify these motifs, we lowered the criterion of preservation in the family, but this is not always necessary. We found a family with a double supercoiled motif, where both ends wrap around the lasso loop three and four times (LS_4_LS_3_), families with a loop pierced four times (L_4_), family where the lasso is embedded inside the knot (L_1_K3_1_), and an example of how both the lasso and the knot topology can exist on a single chain (L_3_#K3_1_). The existence of lasso-knot topologies provides clues on how such structures can be folded.

To further ensure that these topologies are well predicted, for all of the new topological categories, a sample of five proteins was taken and recomputed using the AlphaFold 3 prediction model [34], and analyzed using the AlphaLasso web server. The models proved to have the same topology as those predicted using AlphaFold 2. Below we characterize each of the new motifs in detail.

### *The new lasso topology – L*_4_

We identified two groups of proteins with a new L_4_ lasso topology – L_+4C_ and L_-4C_ (Figure 4). In AlphaLasso we identified 1,082 proteins possessing the L_+_4C lasso and InterPro family ID IPR014635. This number corresponds to 82% of all proteins analyzed for this InterPro family, 61% of its entirety. Evidence suggests that such topology is strongly conserved in this family. On top of that, we found 1,249 proteins with L_-4C_ and InterPro family ID IPR043538, which correspond to 23% of the analyzed proteins with the same InterPro family. While this ratio is rather small in comparison to the first family, it is still visible as an outlier in our compiled data. These two families are not the only ones in which we found L_4_ structures, however, other families did not contain such high numbers of lassoed proteins or lacked a significant representation in the InterPro database.

**Figure 4:**
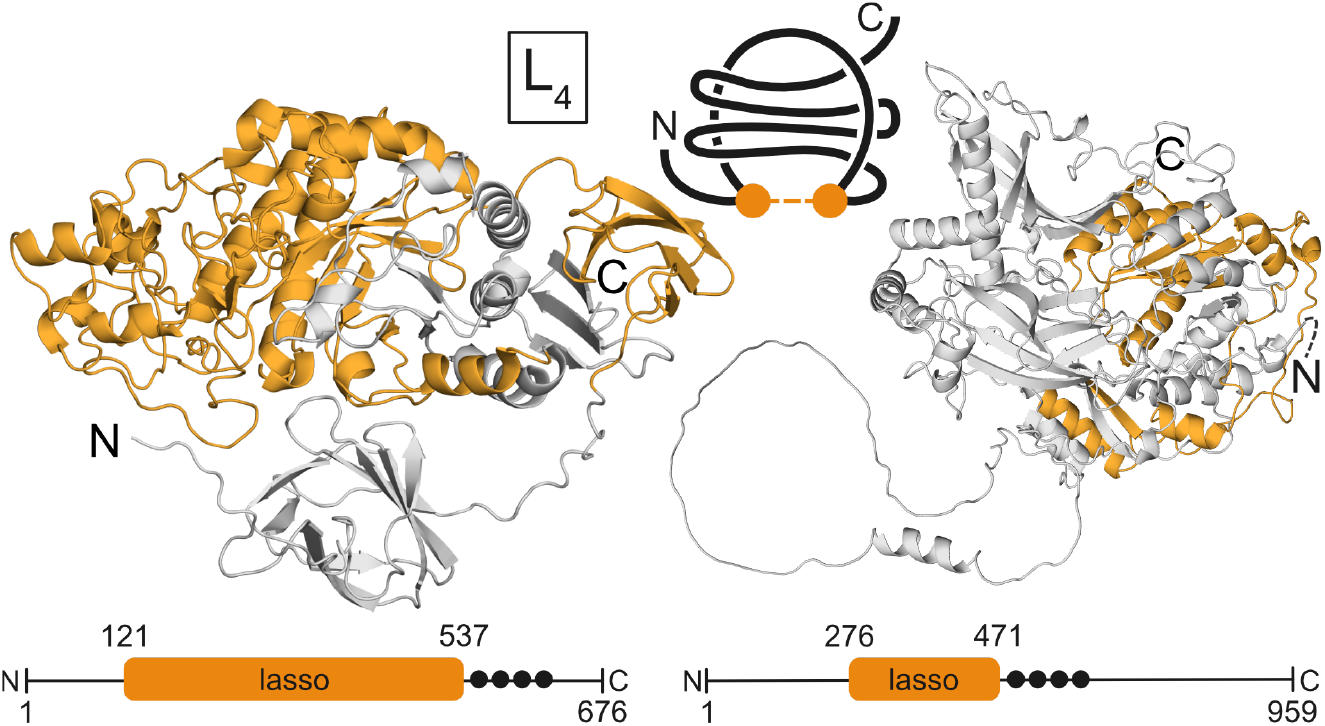
Proteins with the new L_4_ lasso type. Left: L_+4*C*_ lasso in Alpha-Amylase protein. Cartoon and schematic representation of AlphaFold model of *Escherichia coli* alpha-amylase (UniProtKB id: P25718). The orange color shows the position of the lasso loop. The orange bead and line connecting them depict a cysteine bridge that forms the lasso loop. The lower panel shows the position of the lasso motif. The black dots mark the four residues that pierce the lasso loop (Tyr547, Ala564, Gln578, Phe623). Right: L _-4*C*_ lasso in Xylosyltransferase 1 protein from *Homo sapiens* (UniProtKB id: Q86Y38). The first 142 residues are not shown in the figure (schematically shown with a dashed line). The black dots mark the four residues that pierce the lasso loop (Gly491, Gln519, Thr534, Met545).

**L**_+**4C**_ **in Alpha-Amylase Proteins** The L_+4C_ lasso type protein identified with InterPro ID IPR014635 belongs to the family 13 (GH13) of the glycosyl hydrolases from alpha-amylase proteins, which consists of more than 20 types of enzymes [35]. They can be found in a variety of taxonomic groups, like Archaea, Bacteria, Fungi, Plants, and Animals [36, 37, 38]. However, all proteins with L_+4C_ identified belong to Bacteria. Enzymes from this family act on starch and starch derivatives with hydrolysis or transferring activity [39]. Most alpha-amylases consist of three domains, an N-domain, a catalytic domain, and a C-domain, whereas conventional small amylases only have a catalytic domain and a C-domain. The L_+4C_ proteins found in AlphaLasso present an N-terminal extension and instead of three, there are four domains (N’-domain, N-domain, catalytic domain, and a C-domain). The overall structures share similarities with the first two reported proteins with the same conformation, *Pyrococcus furiosus* thermostable amylase (PFTA), a hyperthermophilic amylase isolated from the archaeon *Pyrococcus furiosus* [40] and Archaic Hyperthermophilic Maltogenic Amylase (SMMA) from *Staphylothermus marinus* [41]. Both proteins, PFTA and SMMA, display four domains, with an N’-domain containing ten and nine beta-sheets for PFTA and SMMA, respectively.

As a representative protein for the alpha-amylase family, we chose a *Cronobacter turicensis* alpha-amylase (UniProtKB ID: C9Y338) with global pLDDT equal to 93.3 (Figure 4). The protein has 663 residues and the cysteine bridge is located between residues 108 and 524, with loop length equal to 417 residues. The average pLDDT for the residues within the lasso corresponds to 95.7. Figure 4 exemplifies the L_4_ topology identified in the Alpha-amylase, in which the loop region formed by the S-S bridge is pierced four times.

#### L_-4C_ in Xylosyltransferase

The InterPro family ID IPR043538 adopts an L _-4*C*_ lasso type in only 24% of proteins in this family. This protein family is functionally associated with xylosyltransferases. This family of enzymes catalyzes the transfer of xylose residues to specific acceptor molecules, primarily proteins. This process, known as xylosylation, is crucial for biological functions in organisms [42]. The expected taxonomic range for this enzyme is Eukaryota and Bacteria. However, all L_-4C_ lasso proteins identified with ID IPR043538 display only members from Eukaryota. In AlphaLasso, L_-4C_ lasso proteins correspond to xylosyltransferase 1, 2 (XT1 and XT2), and type II transmembrane proteins (single-pass transmembrane protein) consisting of a short amino-terminal region in the cytosol, a single transmembrane helix, a stem region required for Golgi localization, a catalytic GT-A domain, and a unique C-terminal domain of unknown function [43].

The Xylosyltransferase 1 protein, AlphaFold ID A0A0S7LW33 with global pLDDT equal to 96.75, Figure 4, represents the protein family with InterPro ID IPR043538. The protein has 397 residues and the cysteine bridge is located between residues 5 and 200, and the loop length is equal to 417 residues. The average pLDDT for the residues within the lasso corresponds to 95.7.

### Knotted proteins with lasso topology

We took our analysis further and cross-referenced the proteins containing a lasso with those containing a knot. The knotted topology is observed substantially less frequently than the lasso (1 out of 250 proteins is knotted [22]), therefore, we did not expect a significant overlap. We found two families with a high content of proteins with the mixed lasso-knot topology. Both families have the simplest 3_1_ knot, the most frequent knot type in the protein world. They differ in the complexity of the lasso topology, measured by the number of lasso piercings – L_1_ and L_3_. Interestingly, both of these families represent membrane proteins.

#### Lasso intertwined with a knot – L_1_K3_1_

Sodium/calcium exchanger protein family is a known group of knotted membrane proteins [44, 33]. Even though the presence of the 3_1_ knot in these proteins is known, the fact that the majority of them can also simultaneously form a lasso topology has been unrecognized until now. In particular, we found that 2,952 proteins (65% of the analyzed proteins in the family; based on AlphaFold models of the InterPro family with id: IPR004836) form a single pierced loop closed via a cysteine bridge (L_+1C_ type).

Figure 5 shows the example SLC8 human protein. It has 3_1_ knot spanned on the majority of its multi-domain structure. The knot starts and ends in one of the domains – the transmembrane core domain, common for all the proteins of the family. The lasso loop in the SLC8 is made by the cysteine bridge (Cys55-Cys827) and is pierced by the remaining C-terminal chain (Val855 is the piercing residue; it also lies within the knotted region). Given the position of the knot and the lasso in the structure, the topologies are structurally connected and intertwined (Figure 5). Moreover, the remaining tails (the residues outside either of the entangled motifs) are over 50 amino acids long, thus the motifs are deeply embedded in the structure, which indicates its topological stability. Importantly, the same topologies can be found in the experimental structures of this protein (e.g. PDB id: 8sgj [45]).

**Figure 5:**
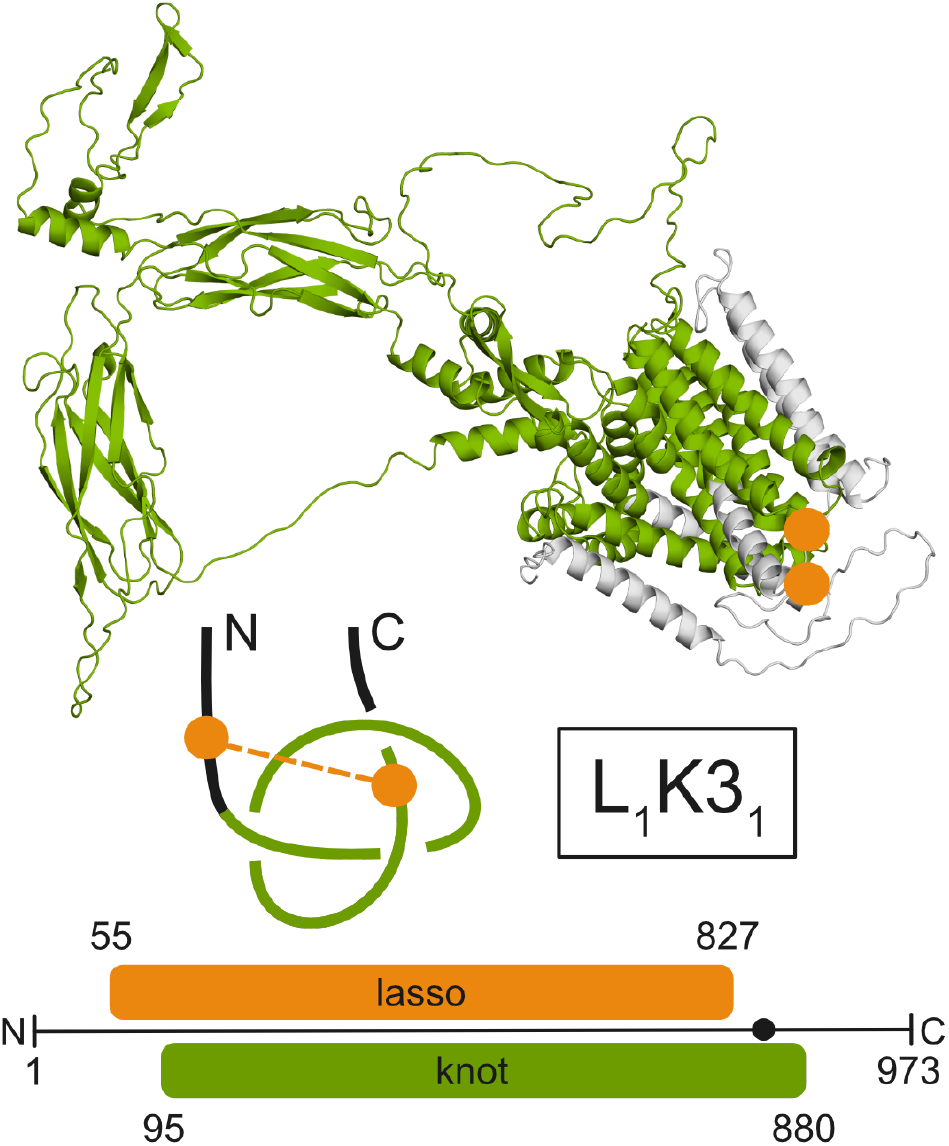
L_+1C_ lasso intertwined with 3_1_ knot in a sodium/calcium exchanger protein. Cartoon and schematic representation of AlphaFold model of the human Sodium/calcium exchanger 1 protein (UniProtKB id: P32418). The green color shows the position of the knot. Orange beads and lines connecting them depict a cysteine bridge that forms the lasso loop. The lower part of the figure shows the positions of the entangled motifs in the protein sequence. A black dot marks the position of the residue that pierces the lasso loop (Val855).

#### Lasso and knot as separate motifs in a single structure – L_3_#K3_1_

The second family with lasso-knot topology is calcium-activated potassium channels (InterPro id: IPR047871). In this case, the lasso and the knot are present in separate domains of the protein. The lasso is in the N-terminal part of the protein (the transmembrane domain), and the knot is located after the lasso motif (the cytoplasmatic domain; Figure 6). The lasso is formed by a short loop (Cys79-Cys206; based on UniProtKB id: Q12791). Note that Cys79 is in an unmodeled region in the experimental structure of this protein (e.g. PDB id: 8z3s) and in the AlphaFold model the pLDDT of this residue is low (53.8). The loop is pierced three times by the C-terminal chain (Ala235, Phe253, Trp268) which makes it a L_+3C_ lasso type. The 3_1_ knot is positioned between Cys413 and Leu1046 and thus needs most of the structure to form (the knot is also present in the experimental structure). The knot is not located in the transmembrane domain like in the case of the sodium/calcium exchanger protein family (Figure 6). Note that both of these entangled motifs are positioned deeply in the protein structure, and any thermal fluctuations of the N- or C-tails will not change the topology of the protein.

**Figure 6:**
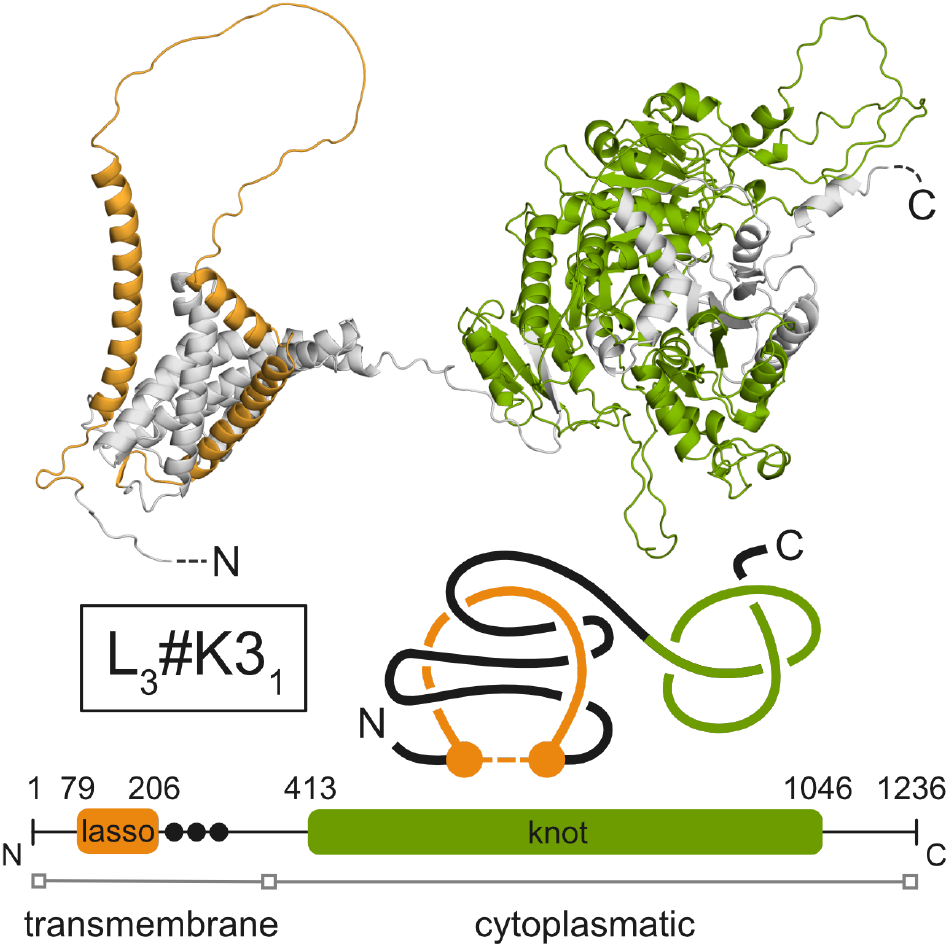
L_+3C_ lasso and 3_1_ knot in a calcium-activated potassium channel. Cartoon and schematic representation of AlphaFold model of the human Calcium-activated potassium channel subunit alpha-1 (UniProtKB id: Q12791). The orange color shows the position of the lasso loop and the green shows the position of the knot. Orange beads and lines connecting them depict a cysteine bridge that forms the lasso loop. The lower part of the figure shows the positions of the entangled motifs in the protein sequence. The black dots mark the position of the three residues that pierce the lasso loop (Ala235, Phe253, Trp268). The positions of the transmembrane and cytoplasmic domains are taken from the UniProtKB.

The lasso-knot topology is not commonly found in this family (31% of the proteins). However, our topological analysis shows that an additional 15% possess knots (but without the lasso). The next 5% only have the lasso topology. Given that the two motifs are found in separate domains, it appears that the overall topology relies on the protein architecture, and in the case of this family, it is not particularly conserved.

### The most complex lasso motif – LS_4_LS_3_

During our extensive search of lasso proteins among AlphaFold models, we encountered a group of proteins with the most complex lasso motif – a double supercoiled lasso with four and three crossings (from N and C-tail, respectively; LS_4_LS_3_; Figure 7). Not only are both tails repeatedly piercing the lasso loop, but they are also wrapped around the chain of the loop (supercoil).

**Figure 7:**
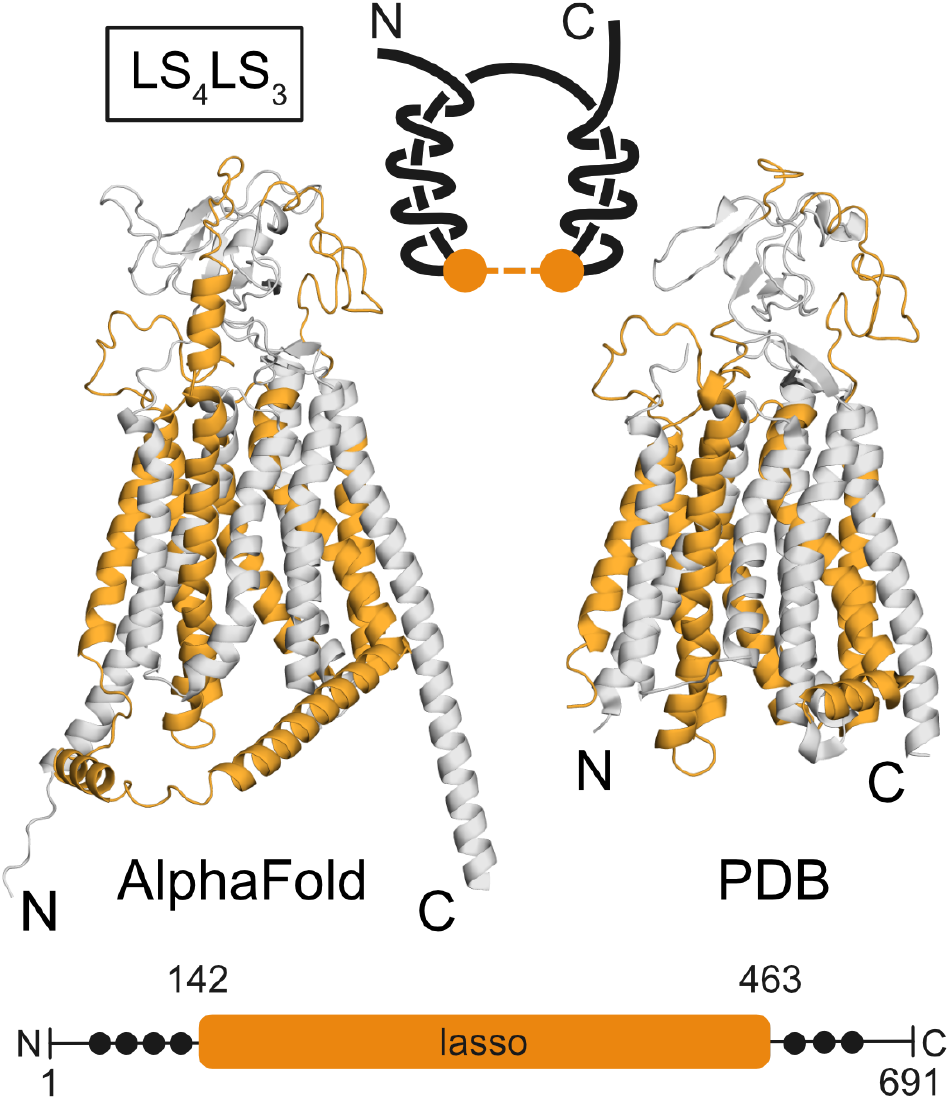
Schematic representation of the protein with the most complex lasso topology (LS_4_LS_3_). Superimposed AlphaFold model (left) and PDB structure (8hnd; right) of Q9Y6L6 protein. The chain of the lasso loop is colored with orange. The black dots mark the position of the residues that pierce the lasso loop (N-tail: Ile47, Phe59, Val80, Arg93, and C-tail: Trp470, Gly552, Ser576).

This complex motif is present in a family of membrane transporters (Organic-Anion-Transporting Polypeptides; InterPro id: IPR004156). The representative protein of this set is human Solute carrier organic anion transporter family member 1B1 (OATP1B1; UniProtKB id: Q9Y6L6), which has structural information available (6 cryo-EM PDB structures). The structures were obtained in different conditions (ligands), leading to their different conformations (outward- and inward-open). We observed that regardless of the conformation, the structures show a complicated lasso topology; however, not as complex as the AlphaFold model (Table S1). The difference lies in the number of times the tails pierce the lasso loop. In particular, the PDB structure 8k6l has one less N-tail crossing than the AlphaFold model (LS_3_LS_3_ vs. LS_4_LS_3_). Both structures are high-quality (the PDB structure 8k6l has 2.9Å resolution and the AlphaFold model has an average pLDDT 80.8) and they superimpose well (C-alpha RMSD 1.5Å based on 320 residues). A closer investigation of the structures shows that the PDB structure lacks a portion of the lasso loop residues (Pro280-Val322). The location of this gap is directly responsible for the missing N-tail crossing.

The lasso loop of OATP1B1 protein is made with a cysteine bridge between Cys142 and Cys463, which are part of different external loops (Cys142: EL3-4 called N-lariat, and Cys463: EL9-10 called Kazal domain). This disulfide bond is present throughout all the structures we analyzed, both experimental (including their different conformations) and theoretical ones (Table S1). The AlphaFold model of the protein is of high quality, including the region with the lasso loop (average pLDDT value 82.6). The model resembles the inward-open conformation of the protein as evidenced by the comparison with the experimental structures (1.7Å with the inward-open conformation (PDB id: 8hnd; Figure 7) and 5.9Å with the outward-open conformation (PDB id: 8k6l), based on 560 and 547 C-alpha atoms, respectively).

### Potential folding pathway

The identification of new types of lasso motifs is also relevant for the fundamental processes of protein folding and degradation. The size (length) of these proteins is too large to conduct quantitative and qualitative simulations of folding even in a coarse-grained model such as Go-like. However, it is possible to propose possible pathways, and we will present one for the most complex lasso motif – LS_4_LS_3_ (Figure 8).

**Figure 8:**
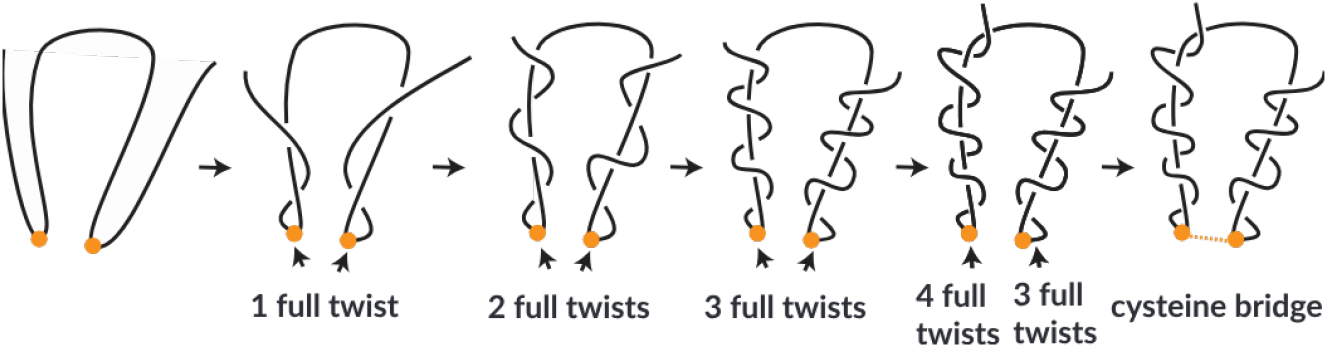
Proposed protein folding mechanism of the most complex lasso topology (LS_4_LS_3_). Positions of the cysteine residues are marked with orange circles.

To the best of our knowledge, for only one protein with a supercoiling (LS_3_) motif, the tangling mechanism has been proposed based on numerical simulation. It was found with structure-based models [46] and molecular dynamics that native contacts are sufficient to self-coil a protein [47] via threading one tail around a closed loop, even when the cysteine bridge was created first.

In the case of LS_4_LS_3_ motif, the only viable option is to wrap the termini around the preformed lasso loop and then close it via a cysteine bridge. The wrapping can be done by a triple or quadruple twisting and then locating the protein in the membrane and closing the loop, see Figure 8.

## Conclusions

Herein, we have shown that Alphafold 2 has the capability of predicting novel lasso topologies. The motifs present in this paper have a significant representation in their InterPro families, suggesting a strong evolutionary conservation. Apart from the *L*_4_ motif, each new lasso is present in a single protein family. The *L*_4_ is found in two enzymatic families: alpha-amylases and xylosyltransferases. Even though they share the same type of lasso topology, the motif itself is resolved differently in each family, e.g. there is a twofold difference in length.

By compiling all predicted 3D structures with lasso and knot topologies, we found two families containing proteins with mixed lasso-knot topologies (L_1_K3_1_ and L_3_#K3_1_). Therefore, for the first time, we show that two complex, entangled motifs can be present in a single structure. Moreover, the motifs are located deeply in the protein structure, indicating their topological stability. Both of these families represent membrane proteins.

The most complex lasso motif (LS_4_LS_3_) is the first two-sided supercoiled lasso found in protein structures. As such, it gives new insight into the protein folding mechanism. We propose a pathway that includes the formation of a supercoiled structure first (by twisting), and then closure by a cysteine bridge.

Based on the newly available data, we show that the lasso topologies are best conserved for loops that are 5-100 amino acids long. By comparing the data generated by the AF to experimentally determined structures available in LassoProt, we confirmed that the AF correctly predicts lasso topologies for already known structures.

The computational approach, such as AlphaFold, has proven advantageous in predicting proteins with unseen-before types of non-trivial topology. Taking into account data presented here, AF can be further used to predict new not-seen-before lasso topology in proteins with new desired properties, so that, for example, they perform functions under unnatural conditions.

## Materials and methods

**Data set** As a data source for our analysis, we utilized AlphaFold-predicted models of protein structures with an average pLDDT≥ 70 to ensure high-quality data. Using the AlphaFold Protein Structure Database (4th version) we generated a list of over 176 million models that met this criterion.

The structural data of AlphaFold-predicted models were analyzed in search of cysteine bridges and lasso topologies [11] during the creation of the AlphaLasso database [30]. The detection of cysteine bridges was based on a distance criterion between sulfur atoms (≤ 3 Å) using Biopython (version 1.79) for processing PDB files. The Topoly (version 1.0.2) [48] Python package was utilized to determine lasso topology. Similarly, such data was processed during the development of the AlphaKnot database [31], which also employs the Topoly Python package for knot detection.

Using the AlphaLasso database, we retrieved proteins with lasso topologies (2.6M individual proteins, 3.1M lassos) along with their corresponding InterPro family IDs and stored them in a PostgreSQL (version 16.2) database. Additionally, data on knotted proteins (681K proteins) were imported from the AlphaKnot database. This integration allowed for efficient querying and aggregation of the data. UniProt IDs were mapped to InterPro family IDs, using the protein2ipr dataset (1.4B pairs) obtained from the InterPro website. Such curated data was then processed using SQL and the pandas (version 2.2.2) package to quantify the proteins within each InterPro family that contained either knots, lassos, or both motifs. For each InterPro family, the ratio of proteins having interesting features to the total (present in InterPro DB) and the analyzed number of proteins was calculated. Only InterPro entries for which we had run the calculations for at least 30% of all the representative proteins were taken into consideration.

Data used in this work is available in the supplementary material. For the data used for discovery of new topologies see S1.1, data used for the comparison with experimental data see S1.2, data used for the analysis of loop size see S1.3.

From each category of topology present in this paper, a sample of 5 proteins was taken and recomputed using the AlphaFold 3 prediction model [34]. The obtained models were then analyzed using the AlphaLasso web server. The models proved to have the same topology as those predicted using AlphaFold 2.

## Supplementary material

*S1*.*1 File* **SI_AlphaLasso.xlsx** – spreadsheet containing information about InterPro families present in the AlphaLasso database. Field summary:

- InterPro ID – entry ID in the InterPro database.
- Type – lasso type.
- Lasso Type (LT) – number of entries of given InterPro ID and Type present in the AlphaLasso database. Represents count of lasso topology. E.g. protein possessing two lassos of L1 type would add 2 to this number.
- Unique Lasso Type (ULT) – number of proteins of given InterPro ID and Type present in the AlphaLasso database. Represents count of proteins possessing at least one lasso of given type. E.g. protein possessing two lassos of L1 type would add 1 to this number.
- Unique in AlphaLasso (AL) – number of proteins of given InterPro ID present in the AlphaLasso database. Includes all types of lassos.
- In InterPro (IP) – number of proteins of given InterPro ID present in InterPro database.
- Analyzed by AL (AI) – number of proteins of given InterPro ID that were analyzed by AlphaLasso web server.
- ULT/IP – ratio of Unique Lasso Type (ULT) to In InterPro (ID). Represents conservation of all known proteins of this InterPro ID.
- ULT/AI – ratio of Unique Lasso Type (ULT) to Analyzed by AL (AI). Represents conservation of all analyzed proteins of this InterPro ID. This metric is used as a conservation value in the paper.
- AI/IP – ratio of Analyzed by AL (AI) to In InterPro (ID). Represents how much of the data available in InterPro was processed by AlphaLasso.

*S1*.*2 File* **SI_Experimental_validation.xlsx** – spreadsheet containing information about InterPro families present in both AlphaLasso and LassoProt databases. Field summary:

- InterPro ID – entry ID in the InterPro database.
- Type – lasso type.
- Unique Lasso Type (ULT) – number of proteins of given InterPro ID and Type present in the AlphaLasso database. Represents count of proteins possessing at least one lasso of given type. E.g. protein possessing two lassos of L1 type would add 1 to this number.
- In InterPro (IP) – number of proteins of given InterPro ID present in InterPro database.
- Analyzed by AL (AI) – number of proteins of given InterPro ID that were analyzed by AlphaLasso web server.
- Conservation – ratio of Unique Lasso Type (ULT) to Analyzed by AL (AI). Represents conservation of all analyzed proteins of this InterPro ID. This metric is used as a conservation value in the paper. Represented as percentage in increments of 10.
- AI/IP – ratio of Analyzed by AL (AI) to In InterPro (ID). Represents how much of the data available in InterPro was processed by AlphaLasso.

*S1*.*3 File* **SI_Loop_size.xlsx** – spreadsheet containing information about loop size distribution in InterPro families present in AlphaLasso database. This data consists only of InterPro entries for which at least 50% (minimum 100) of entries present in the InterPro database were calculated. Field summary:

- InterPro ID – entry ID in the InterPro database.
- Type – lasso type.
- Conservation – ratio of Unique Lasso Type (ULT) to Analyzed by AL (AI). Represents conservation of all analyzed proteins of this InterPro ID. This metric is used as a conservation value in the paper. Represented as percentage in increments of 10.
- Loop size – mean loop size (in amino acids) of all proteins in the given InterPro entry.

*S1 Table* **Detailed topology information about different structures of the protein with the most complex lasso motif (UniProtKB id: Q9Y6L6).**

The summary of available PDB and AlphaFold structures, comparing calculated topology for proteins with different ligands.

## ACKNOWLEDGMENTS

This work was supported by the National Science Centre 2022/47/B/NZ1/03480 (to J.I.S.) This research was carried out with the support of the Interdisciplinary Centre for Mathematical and Computational Modelling (ICM) University of Warsaw under computational allocation no GS82-12.

